# Universality of the developmental origins of diverse functional maps in the visual cortex

**DOI:** 10.1101/354126

**Authors:** Min Song, Jaeson Jang, Gwangsu Kim, Se-Bum Paik

## Abstract

The primary visual cortex of higher mammals is organized into diverse functional maps of correlated topography, implying an efficient tiling of functional domains. However, no fundamental principle is available on how systematic organization of the maps could develop initially in various species. Here, we propose that diverse functional maps are seeded from a common framework of retinal afferents, and that this universality of development explains the observed topographical correlations among the maps. From the simulation of retinal ganglion cell mosaics, we successfully developed all four cortical maps observed so far. We validated our model prediction of inter-map relationships from analysis of map data in evolutionarily divergent mammalian species. Our key prediction, the hexagonal periodicity of every functional map, was validated from analysis of the map data in diverse mammalian species. Our results provide new insight into a universal mechanism for the development and evolution of the visual cortex.

## Introduction

In higher mammals, the primary visual cortex (V1) is organized into various functional maps of neural tuning such as ocular dominance^1^, preferred orientation^2^, direction^3^ and spatial frequency^4^. Although the role of each functional map in visual information processing is under debate^5,6^, correlations between the topographies of different functional maps have been observed, implying their systematic organization. For instance, it was reported that the gradient of orientation tuning intersects orthogonally with that of ocular dominance^7^ and preferred spatial frequency^8^ in the same cortical area. High-resolution two-photon imaging data revealed that the region of higher spatial frequency tuning tends to align with the binocular region in the ocular dominance map^9^. Such structural correlation between the maps is thought to achieve a uniform representation of visual features across cortical areas^10^. In addition, these findings in phylogenetically distinct mammalian species imply that there may exist a universal principle of development and evolution organizing individual functional maps^11^, possibly which results in efficient tiling of sensory modules.

Important clues regarding the development of the maps were found in the thalamic origin of the selective response in V1. When cortical recurrent activity was silenced so that only thalamic afferents were provided, the preferred orientation angle of a V1 neuron remained consistent^12,13^, which suggests that cortical orientation tuning originates from feedforward afferents. In addition, it was reported that orientation tuning in V1 is predictable from the local average of thalamic ON and OFF receptive fields^14^. At larger scales, it was reported that the topography of functional maps in V1 is strongly correlated with the spatial arrangement of thalamic afferents^15,16^. Furthermore, it was recently observed that direction preference in V1 arises from temporal delay in thalamic inputs^17^. These observations altogether imply that functional tuning in the cortical neurons may originate from a common retinal or thalamic afferent (**Fig. 1a**); thus map structures of various functional tuning might be seeded initially from the same blueprint of feedforward projections.

**Figure 1.**
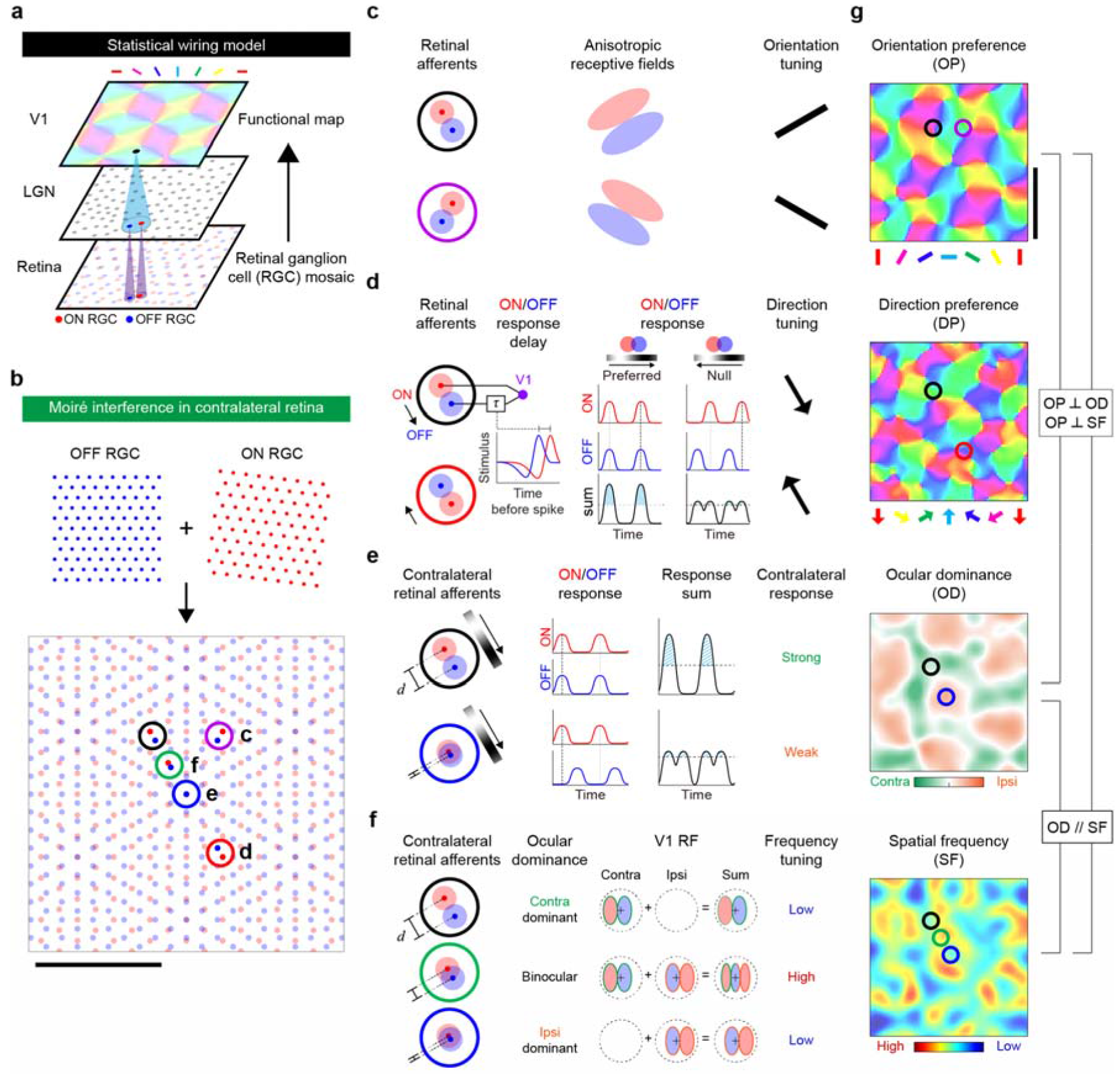
Retinal origin of various functional maps in V1. **a**, Developmental model of cortical functional maps as a feedforward projection of ON and OFF retinal ganglion cell (RGC) mosaics. **b**, The moiré interference pattern between ON and OFF RGC mosaics in the contralateral retina provides a common framework of hexagonal patterns for cortical maps. The scale bar indicates the period of the interference pattern. **c**, With the local feedforward pooling of RGCs, orientation tuning of V1 neurons is induced from the alignment of ON and OFF subregions. **d**, Temporal delay between ON and OFF pathways result in a direction-selective response of V1 neurons. Preferred direction of the neuron is from ON to OFF centres of RGCs. **e**, According to the distance between neighbouring RGCs in the contralateral retina, the degree of overlap between ON and OFF subregions in V1 receptive fields varies and differential response occurs. When ON and Off subregions are far from each other, the V1 neuron can generate a strong response to contralateral input. If ON and OFF RGCs are very close; they are not simultaneously activated and cannot respond strongly to contralateral inputs. **f**, In binocular regions, phase difference between the contra- and ipsilateral receptive field induces preference for higher spatial frequency. **g**, The common retinal origin of each functional map causes a topographical correlation among the maps. The scale bar indicates the period of the orientation map, which is set up by the interference pattern in **b**.

Given this, how could the feedforward afferents provide a regularly structured layout of functional tuning to seed an orderly architecture of the map? The statistical wiring model^18,19^ suggested that V1 receptive field structure is constrained by the local structure of ON and OFF mosaics of retinal ganglion cells (RGCs)^20^. In this view, the receptive field of a V1 neuron is generated from the sum of receptive fields of local RGC that provides feedforward afferents, and the V1 neuron is tuned to orientation due to the anisotropy of the receptive fields. Adapting this notion, the moiré interference model further proposed that spatial interference patterns between the hexagonal mosaics of ON and OFF RGC generate a quasi-periodic pattern that seeds the topography of an orientation map early in development^21,22^ (**Fig. 1b-c**). The key assumption that RGC mosaics patterns seed the topography of cortical orientation maps, allowed this model to successfully explain how regularly structured cell mosaics in the retina could seed the early structure of an orientation map^23^ and its hexagonal periodicity^24^ in diverse species of higher mammals.

Herein, expanding this developmental principle of orientation maps, we propose that such a regular distribution of RGCs can also induce multi-modal functional maps in V1 (**Fig. 1d-g**). Our hypothesis is that local ON and OFF afferents can induce multiple types of neural tuning according to their spatial and temporal profiles. First, temporal delay between ON and OFF pathways induces direction selectivity in receptive fields of ON and OFF subregions (**Fig. 1d**). Second, variation of the distance between neighbouring ON and OFF RGCs modulates the degree of overlap between ON and OFF receptive field subregions, which varies the sum of neural responses to ON and OFF stimulus in contralateral afferents. This induces competition between contra- and ipsilateral afferents, resulting in a periodic pattern of ocular dominance (**Fig. 1e**). Lastly, in the binocular region, intermingling of both contra- and ipsilateral inputs with different spatial phase of receptive field induces a preference to higher spatial frequency than that in the monocular region (**Fig. 1f**). Based on this model, simulations were able to reconstruct successfully each functional map and the observed topographical correlations between them (**Fig. 1g**).

We validated our model predictions using published animal data on evolutionarily divergent species^7,10,25–30^. We confirmed that all the predicted relationships between the local structures of different functional maps from the model were observed in experimental data. In addition, our new prediction that iso-domains of feature preference are arranged in a hexagonal lattice in each type of functional map was confirmed using map data from various species. Taken together, we suggest that the regularly structured retinal afferents provide a common framework of organizing various functional maps in the visual cortex.

## Results

We propose a new theory that the layouts of cortical functional maps are seeded from the spatial distribution of ON and OFF ganglion cells in the retinal mosaics (**Fig. 1**). To validate this model, first we performed a series of computer simulations using an ideal model of retinal mosaics where the locations of ON and OFF receptive fields lie at the vertices of hexagonal lattices^31^ (**Fig. 1b**). In our previous study^21^, the model showed how a single pair of ON and OFF RGC mosaics from the contra-lateral eye could initially generate the structure of orientation maps^23^ (**Fig. 1c**). In the current study, we expand this model to consider both contra- and ipsilateral afferents to V1, to investigate the emergence of other functional maps. Our results show that retinal afferents could induce periodic patterns of direction preference (**Fig. 1d, Fig. 2**), ocular dominance (**Fig. 1e, Fig. 3**) and spatial frequency preference (**Fig. 1f, Fig. 4**), from the regularly structured moiré interference pattern of RGC mosaics.

**Figure 2.**
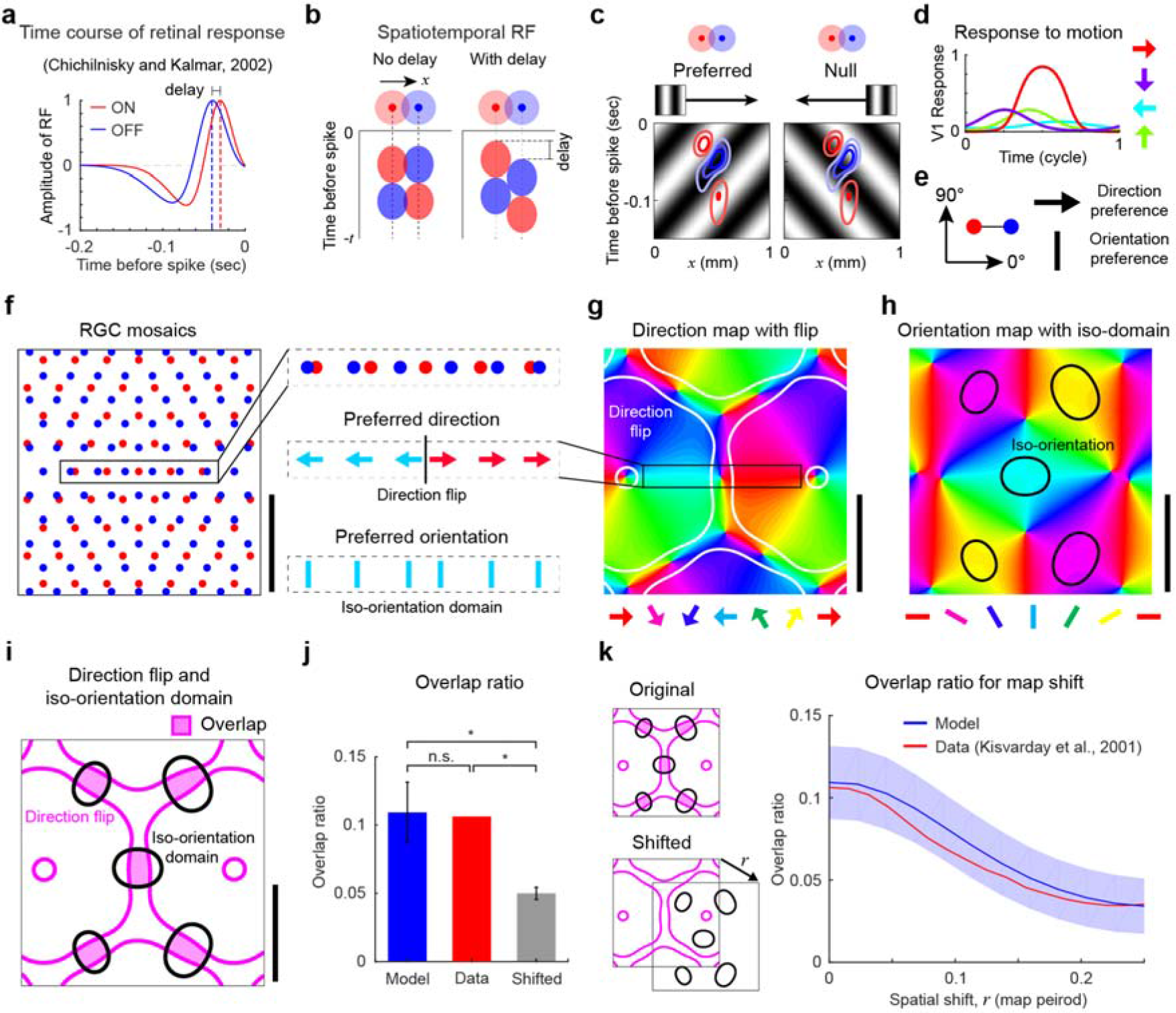
Flip of preferred direction at the centre of iso-orientation domains. **a-e**, Direction tuning of a V1 neuron originates from a temporal delay between ON and OFF pathways. **a**, Temporal dynamics of receptive fields indicates a consistent delay (~9 ms) between ON and OFF pathways^32^. **b**, The ON-OFF delay induces a non-separable spatiotemporal receptive field of V1 neuron. **c**, This non-separable receptive field induces direction tuning with preferred response to ON-to-OFF direction. **d**, Direction tuning of a model V1 neuron. **e**, Preferred direction and orientation from an ON-OFF dipole structure. **f**, In moiré interference, the direction of ON-to-OFF RGC dipoles suddenly flips in areas where their orientation is consistent. The scale bar indicates the half period of the interference pattern. **g-i**, Direction and orientation maps seeded from a common origin of the RGC mosaics in **f**. The scale bar indicates a half-of-map period. **g**, Linear flips (white solid lines) in a direction selectivity map**. h**, Iso-orientation domains (black solid lines) in an orientation map. **i**, Direction flips in **g** and iso-orientation domains in **h**. Direction flip (pink solid lines) appears at the centre of the iso-orientation domains (black solid lines). **j**, Both in model (*N* = 100) and animal data (*N* = 1)^27^, the overlap between the two regions is significant, but not in randomly shifted data (*N* = 100). (model vs. shifted: two-tailed t-test, *P* < 10^−15^, *T* = −25.723; data vs. shifted: one sample two-tailed t-test, *P* < 10^−15^, *T* = −126.146; model vs. data: one sample two-tailed t-test, n.s., *P* = 0.177, *T* = 1.36) Error bars indicate SD. **k**, The average of overlap ratio in the radial direction shows a local maximum at the origin and gradually decreases as one map is spatially shifted, in both model and data^27^. Shaded region indicates SD.

**Figure 3.**
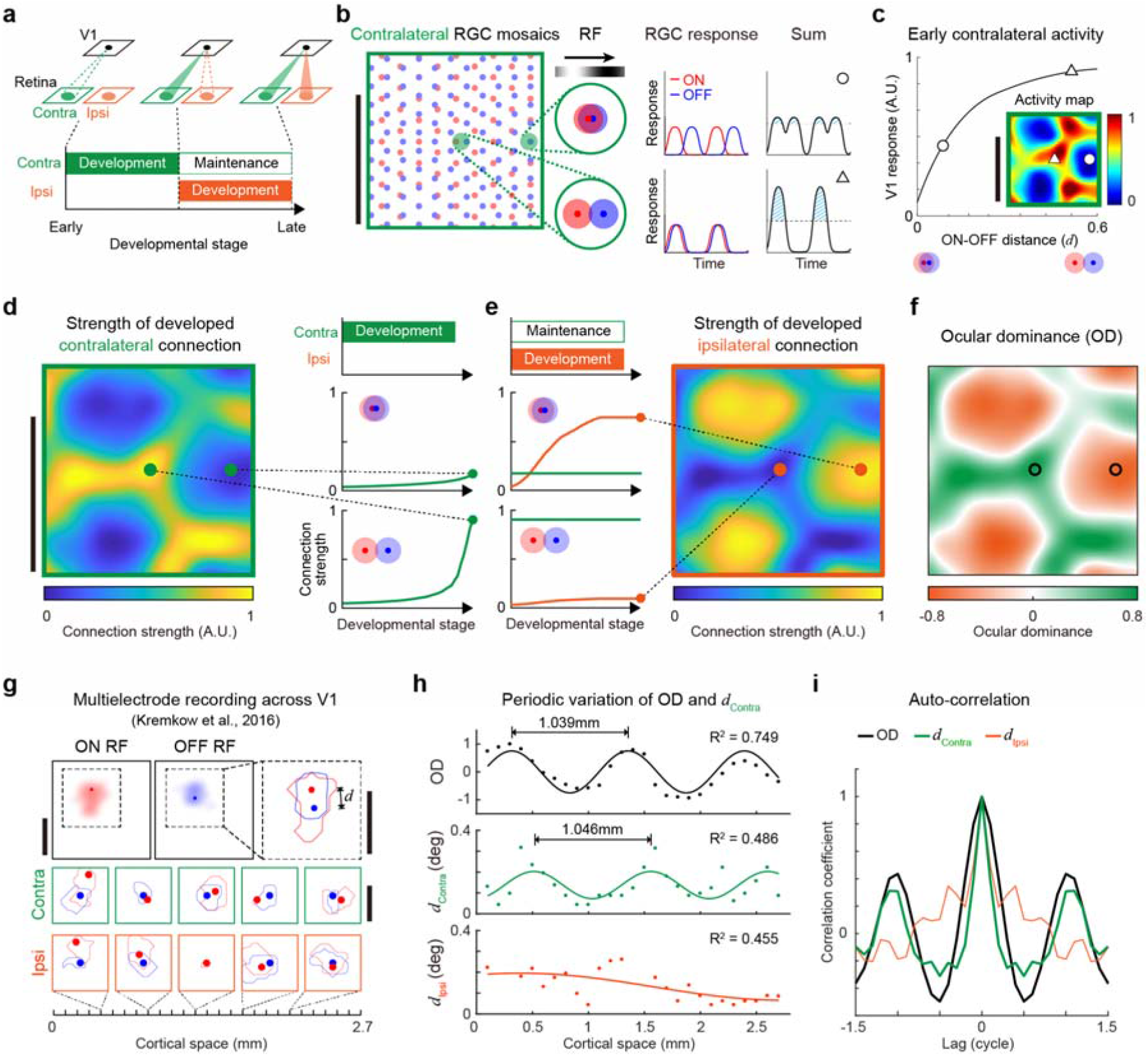
Ocular dominance map from the contralateral retinal mosaics. **a**, Sequential development model of contra- and ipsilateral visual pathways and ocular dominance^33^. **b**, Distance between the neighbouring ON and OFF RGCs determines the degree of overlap in V1 receptive field subregions, and the cortical response. (Top) When two subregions greatly overlap, the neuron induces a weak contralateral response. (Bottom) Where the two subregions show less overlap, the V1 neuron generates stronger response. The scale bar indicates the period of the interference pattern. **c**, The simulated cortical response is an increasing function of the ON-OFF RGC distance. Inset, an early periodic pattern of the cortical response generated by contralateral RGC mosaics. **d**, (Left) Early periodic pattern of the cortical response in **c** induces a periodic pattern of contralateral dominance. (Right) Refinement of the contralateral feedforward connections. An activity-dependent process strengthens feedforward connection when postsynaptic V1 neurons strongly respond. The scale bar indicates a map period. **e**, Refinement of the ipsilateral connections. Strength of total connections from contra- and ipsilateral afferents to a V1 neuron is assumed to be a constant. **f**, Ocular dominance map originated by **d** and **e**. **g**, Previous observation^28^ of receptive field and ocular dominance recorded across the V1 using multiple electrodes. The scale bar indicates 0.5° in visual space. **h**, Both ocular dominance and contralateral ON-OFF distance appeared to vary periodically with the same spatial period (~1 mm), and were fitted to a sinusoidal curve (OD: *R*^2^ = 0.749, contralateral: *R*^2^ = 0.486, ipsilateral: *R*^2^ = 0.455). **i**, Auto-correlation of ocular dominance and ON-OFF distance.

**Figure 4.**
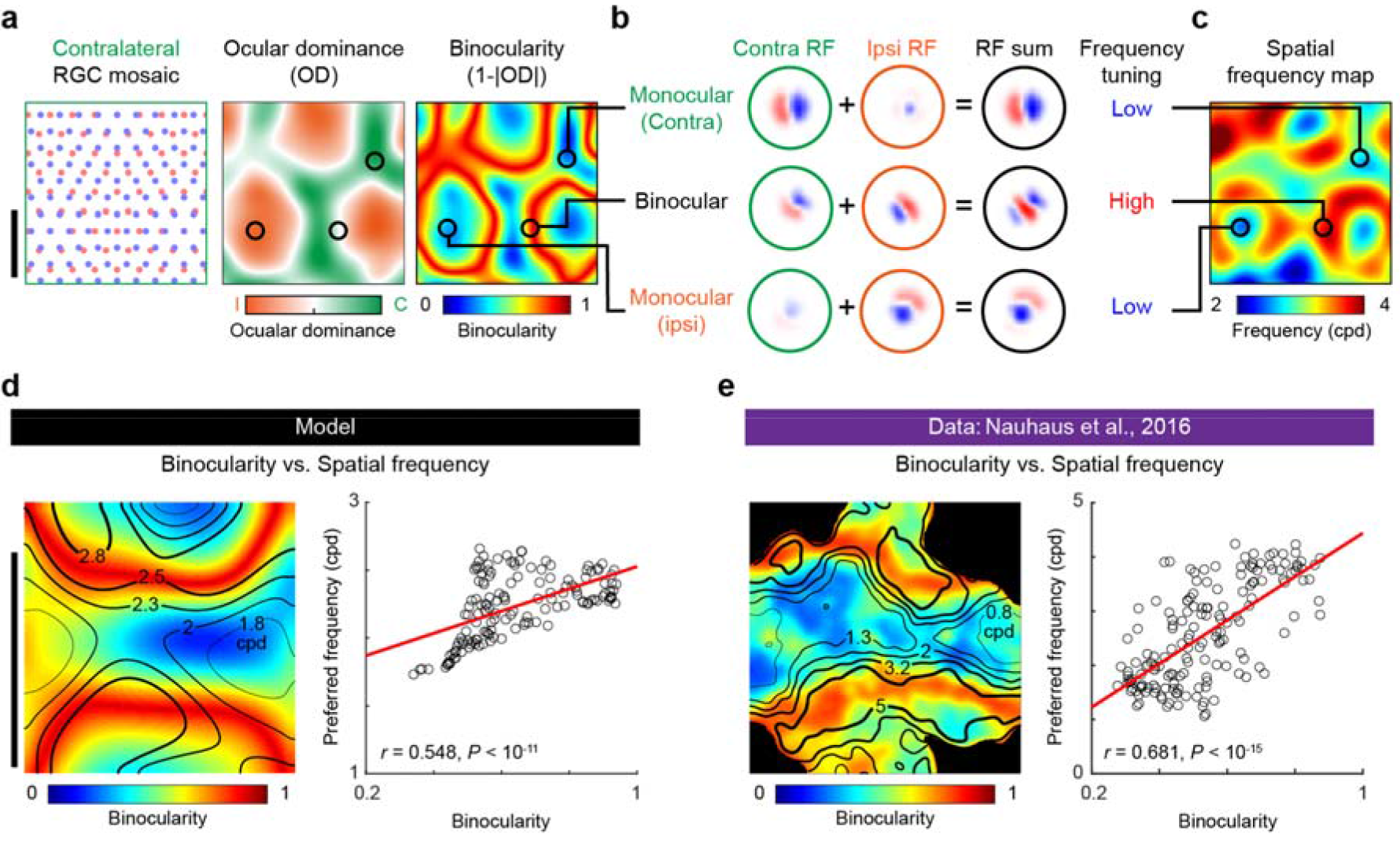
Spatial frequency map arises from the matching of contra- and ipsilateral afferents. **a**, Model ocular dominance map and its spatial organization of binocularity. **b**, Intermingling of contra- and ipsilateral receptive fields in monocular and binocular regions. Phase difference between contra- and ipsilateral receptive fields in the binocular region induces narrower subregions of receptive field, resulting in tuning for higher spatial frequency. **c**, A model spatial frequency map induced from the RGC mosaics in **a**. **d-e**, Simulated (**d**) and observed^9^ (**e**) binocularity vs. spatial frequency maps. (Left) Binocularity map with contours of spatial frequency maps. Thick contours indicate regions encoding high spatial frequency. (Right) A strong correlation between binocularity and spatial frequency maps was observed both in the model and data (Pearson correlation coefficient, model: *N* = 144, *r* = 0.548, *P* = 1.082×10^−12^, data: *N* = 164, *r* = 0.681, *P* < 10^−15^). The scale bar indicates a half period of the interference pattern or map.

### Direction tuning discontinuity in the iso-orientation tuning domain

If the local alignment of ON and OFF retinal inputs constrains orientation tuning (**Fig. 1c**), how does direction tuning arise from these retinal afferents? We hypothesized that difference in temporal response profiles between ON and OFF RGCs could induce the direction tuning of V1 neurons. From the observed delay of response kernels between ON and OFF RGCs afferents^32^ (**Fig. 2a**), a V1 neuron connected to them would have a spatiotemporally non-separable receptive field (**Fig. 2b**), and would induce direction tuning (**Fig. 2c**). This asymmetric spatiotemporal receptive field generates a selective response to the stimulus that drifts from ON to OFF RGCs (**Fig. 2d**), resulting in a preferred direction angle in the ON-to-OFF direction (**Fig. 2e**).

Such common origin of orientation and direction tuning, explains the topographical correlation between the orientation and direction maps. The iso-orientation domains of an orientation map exhibit substantial overlap with the linear fractures of a direction map, in which the preferred direction suddenly flips to the opposite direction^27^. From the notion that the preferred direction of a V1 neuron is constrained by the direction of the ON-to-OFF subregion of a receptive field (**Fig. 2e**), our model predicts that a moiré interference pattern of ON and OFF RGC mosaics contains linear fractures of ON-OFF direction, forming preferred direction discontinuities (**Fig. 2f**). In this view, direction discontinuities would appear in the middle of the iso-orientation domain (**Fig. 2g-i**). To validate this model prediction, we examined previously reported map data of cats^27^, to see if the iso-orientation domains of orientation maps overlap with the direction fractures of the direction map. We confirmed that the overlap ratio observed in the map data was significantly greater than chance (**Fig. 2j**, model: 0.109±0.022, *N*=100, data: 0.106, *N*=1, randomly shifted data: 0.05±0.005, *N*=100; model vs. shifted: *P* < 10^−15^, *T* = −25.723, two-tailed t-test; data vs. shifted: *P* < 10^−15^, *T* = −126.15, one sample two-tailed t-test; model vs. data: n.s., *P* = 0.177, *T* = 1.36, one sample two-tailed t-test), suggesting that orientation and direction maps share a common substrate of spatial organization. On the other hand, when we spatially shifted the direction map from the orientation map and measured again, the overlap ratio noticeably decreased as the amount of shift increased, confirming that the predicted correlation between the two maps does exist (**Fig. 2k**).

### Development of ocular dominance with RGC mosaics from two eyes

We showed that a pair of ON and OFF RGC mosaics (the retinal afferents from the contralateral eye) could seed the layout of orientation and direction maps. However, to describe the development of ocular dominance, the inputs from the ipsilateral eye must be considered as well. With a notion that thalamocortical projections from the contralateral eye arrive at V1 ahead of those from the ipsilateral pathway^33^, we implemented this process so that the connections from each pathway would develop sequentially (Fig. 3a). From the simulation, we propose that the RGC mosaics of the contralateral retina also initiate periodic topography of cortical response by contra- and ipsilateral afferents, and result in the ocular dominance map.

At early developmental stage when only the contralateral afferents are provided, spatial variation of the ON-OFF distance in RGC mosaics could induce periodic fluctuation of local cortical activity (**Fig. 3b-c**). When ON and OFF RGCs are close to each other, ON and OFF subregions in the V1 receptive field largely overlap and cannot be simultaneously activated by visual stimulus. In this case, the cortical responses from ON and OFF afferents are not summed up at their peak values, resulting in weak total response (**Fig. 3b**, top, **Fig. 3c**, white circle). In contrast, when ON and OFF subregions are more separated, both subregions can be simultaneously activated by proper stimuli, resulting in stronger cortical response (**Fig. 3b**, bottom, **Fig. 3c**, white triangle). The estimated cortical response appeared as an increasing function of the distance between ON and OFF RGCs. Moreover, the moiré pattern of contralateral RGC mosaics could induce periodic cortical activity, even without spatial variation in the feedforward connection strengths (**Fig. 3c**). Such a pattern of cortical response at the early stage could provide a blueprint for the ocular dominance map (**Fig. 3d-f**).

To simulate a complete developmental process for an ocular dominance map, we first modelled activity-dependent wiring of the contralateral pathway^33^ such that the strength of feedforward connection was potentiated as the response of a postsynaptic neuron to feedforward spikes got stronger. (**Fig. 3a**; for details, see Methods). As a result, a two-dimensional hexagonal pattern of contralateral wiring strength was developed in the cortex from the ON and OFF RGC mosaics structure (**Fig. 3d** and **Supplementary Fig. 1**). Next, we allowed the same type of plasticity in the ipsilateral pathway and simulated the development of cortical afferents from both eyes. To examine the balance between contra-and ipsilateral afferents at unit scale, we normalized the total strength of contra- and ipsilateral afferents for each V1 neuron as a constant. Thus neurons with strong contralateral afferents received relatively weaker ipsilateral inputs and vice versa (**Fig. 3e**). As a result, this developmental process reproduced a periodic ocular dominance map, the layout of which was initially seeded by contralateral RGC mosaics (**Fig. 3f**).

A key prediction of our model is that the ocular dominance at a cortical site depends on the distance between ON and OFF receptive fields from the contralateral pathway. We were able to validate this prediction using recently published cat data of multi-electrode recordings across the V1^15^ (**Fig. 3g**). We confirmed that the distance between ON and OFF afferents from the contralateral pathway, as well as the ocular dominance, varied periodically across the cortical surface and that their spatial periods were practically identical (only 0.7% different; **Fig. 3h** and **i**). In addition, the observed ocular dominance was significantly correlated with the distance between ON and OFF receptive fields from the contralateral pathway, but not with that from the ipsilateral pathway (**Fig. 3h,** ocular dominance vs. *d*_contra_: *r* = +0.602, *P* = 0.002, lag = 0.2 cycle of ocular dominance period, ocular dominance vs. *d*_ipsi_: *r* = −0.441, *P* = 0.605, lag = −0.5 cycle). This result is consistent with our model prediction that the layout of the ocular dominance map is constrained by the spatial organization of contralateral afferents.

One might argue that the ocular dominance map in a monkey shows a stripe pattern, different from the blob-like pattern in cats and our simulation (**Fig. 3f**). The degree of anisotropy in the retinotopy may provide a clue to explain this difference across species. The focus of our simulation was on the condition of isotropic retinotopy, where magnification factors along the polar and eccentric axes are similar to each other, as observed in the cat^34^. However, in monkeys, the ratio between these two magnification factors is about 1.5^35^, implying that the same distance on each axis in the retinal space is noticeably different in the cortical space. When this anisotropy is introduced in our model, it distorted the initial blob-like pattern, so that blobs were connected along the polar axis, resulting in the stripe pattern along the eccentric axis observed in monkeys (**Supplementary Fig. 2**).

### Development of spatial frequency map from binocular afferents

Next, we show that the ocular dominance patterns in the V1 can induce the organization of spatial frequency maps. In the previous study, it was reported that the spatial organization of frequency tuning in V1 neurons matches the structure of the ocular dominance map in a way that the neurons of higher binocularity (receives strong inputs from both eyes) are tuned to higher spatial frequency^9^. Here, we show that this relationship between spatial frequency tuning and binocularity is achieved from a common origin of the two maps, the spatial organization of ON and OFF afferents (**Fig. 4**).

In our model of ocular dominance maps, local cortical neurons become either a contra- or ipsilateral dominant region, depending on the distance between ON and OFF (**Fig. 4a**). Thus, when the ON-OFF distance in the contralateral RGC mosaics is minimum and maximum, the corresponding V1 regions become monocular (either contra- or ipsilateral dominant; **Fig. 4b**, top and bottom), while the areas in between become binocular regions where the neurons receive balanced input from both eyes (**Fig. 4b**, middle). During this process, we first found that the orientation preference of ipsilateral afferent can match that of contralateral receptive field through an activity-dependent matching process (**Supplementary Fig. 1**), as observed in animal data^23^. More importantly, we found that the phase difference between the contra- and ipsilateral receptive fields^36^ induces a preference to higher spatial frequency of the summed receptive field in binocular regions (**Fig. 4b**, middle). As a result, a periodic spatial frequency map is induced along with the ocular dominance map that is initially seeded from the contralateral retinal mosaics (**Fig. 4c**). From our model simulations, we observed that the spatial frequency tuning was strongly correlated with the binocularity in all local cortical areas (**Fig. 4d**, Pearson correlation coefficient, *N* = 144, *r* = +0.548, *P* = 1.082×10^−12^). Again, we validated this model prediction from analysis of the monkey data used previously^9^ (**Fig. 4e**, Pearson correlation coefficient, *N* = 164, *r* = +0.681, *P* < 10^−15^).

### Topographical correlations of orientation, direction, spatial frequency and ocular dominance maps

How can the observed correlations arise among the diverse functional maps in the cortex? Here, we propose that the structure of retinal afferents provides a blueprint for every functional map in the cortex, and this common origin of RGC mosaics induces intrinsic structural correlations among the maps (**Fig. 5a-d**).

**Figure 5.**
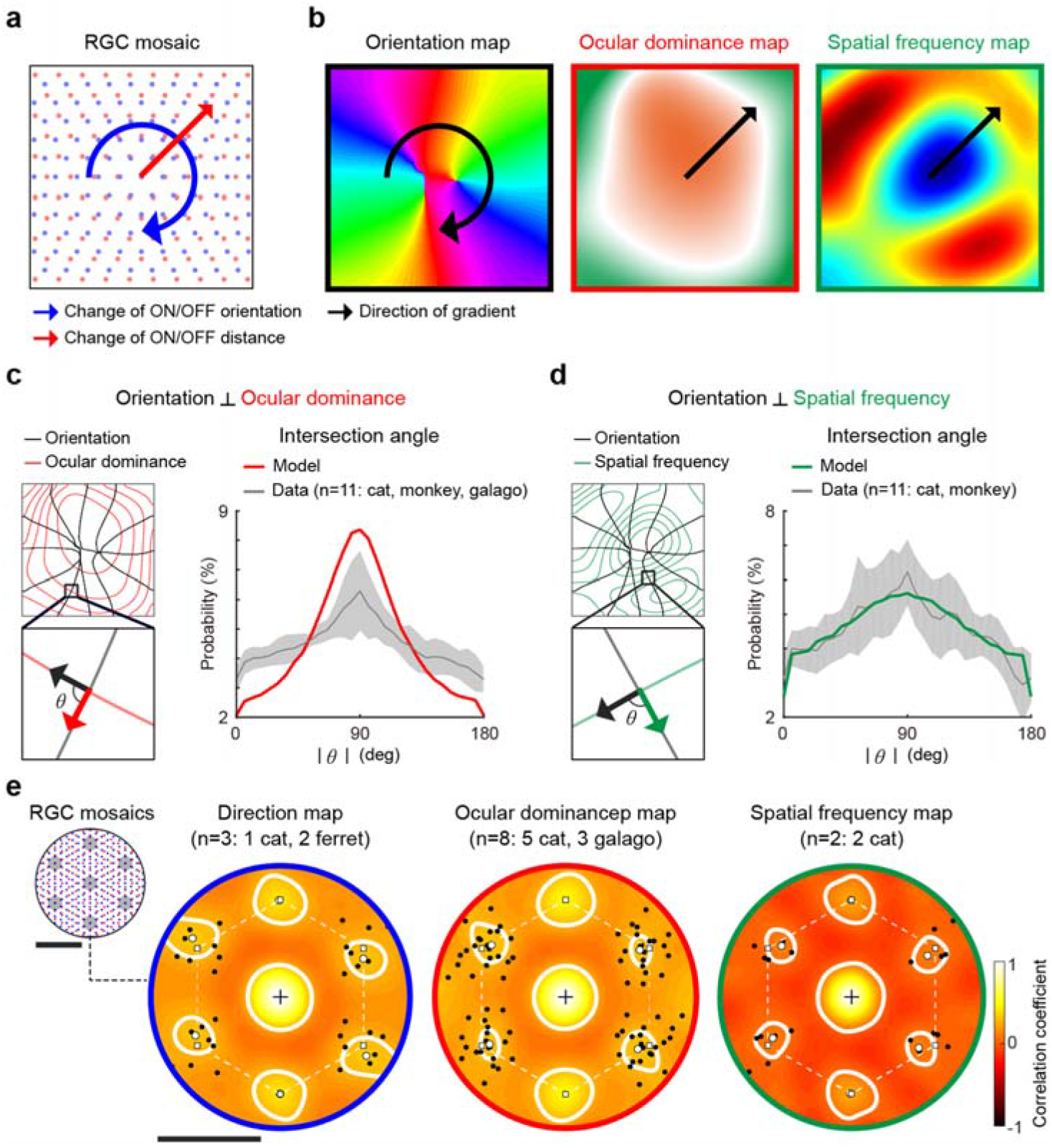
Topographical relationship and hexagonal periodicity of functional maps. **a**, Spatial organization of ON-OFF RGC dipole distance and angular alignment in the moire interference pattern. Note that the gradient of the two variables is orthogonal across the mosaics area. **b**, Orientation preference changes in a circular direction around singularities, while ocular dominance and spatial frequency tunings change in a radial direction. **c**, Orthogonal distribution of intersection angle between the gradients of orientation and ocular dominance maps (11 pairs of maps; 6 cats, 4 monkeys, 1 galago). **d**, Orthogonal distribution of intersection angle between the gradients of orientation and spatial frequency maps (11 pairs of maps; 5 cats, 6 monkeys). **e**, The average autocorrelation patterns of various functional maps from different species^7,10,25–30^. Peaks (open white circles) in the autocorrelation closely matched the ones in a perfect hexagonal lattice (open white squares). Solid black circles indicate peaks in the autocorrelation of individual maps. The scale bar indicates a period of the interference pattern or map.

Previously, we showed that orientation pinwheels develop at the locations where the distance between RGC ON and OFF receptive fields is either local maxima or minima^22^ (**Fig. 5a**). If we examine other functional tunings of neurons around such singularities, it is observed that the orientation, ocular dominance and spatial frequency tunings must have well-correlated map structures, each with contribution from the structure of retinal ON and OFF afferents (**Fig. 5b**). As illustrated in the model RGC mosaics (**Fig. 5a**), the distance between the nearest ON and OFF RGC varies in the direction (**Fig. 5a**, **red arrow**) orthogonal to that of ON-OFF dipole angle variation (**Fig. 5a**, **blue arrow**). Thus, the gradient of a preferred orientation map changes in a circular direction around the orientation pinwheels (**Fig. 5b**, left). At the same time, ocular dominance which depends on the distance between ON and OFF RGCs, changes in the radial direction from the singularity (**Fig. 5b**, middle). As a result, gradients of the two maps must cross orthogonally in the cortical space. Similarly, a spatial frequency map that develops parallel to the ocular dominance map is aligned orthogonal to that of the orientation map (**Fig. 5b**, right).

We validated these model predictions using published functional map data from animal experiments and confirmed the relationship between the maps observed in the simulation. As predicted by our model, both ocular dominance (cat^7,10,25,28,37^, monkey^9,38^, galago^26^) and spatial frequency tuning maps (cat^7,10,25^, monkey^8,9^) showed orthogonal relationship to the structure of the orientation map in the same area (**Fig. 5c-d**). Thus, we concluded that our model, in which cortical functional maps develop universally from the moiré interference patterns in RGC mosaics, could provide a plausible explanation for the development of functional maps and their structural correlations.

### Hexagonal symmetry in diverse functional maps

Another key prediction of our developmental model arises from the intrinsic features of RGC moiré interference patterns: hexagonal symmetry. From the fact that ON and OFF RGC mosaics are hexagonal lattices^31^ and thus also are their interference patterns^39^, our model predicts that each functional map that develops from RGC afferents must have hexagonal symmetry in the map topography in common. We tested this prediction using published animal data on direction^27,29,30^, ocular dominance^7,10,25,26,28^ and spatial frequency maps^7,25^ from different species as was done for orientation maps in our previous study^21^.

As predicted, 2D autocorrelation of the individual maps showed a hexagonal pattern of peaks (**Supplementary Fig. 3**). The statistical significance of the local peaks was tested against control maps that matched the spectral power and spatial periodicity of the original map, but with an isotropic amplitude spectrum instead of hexagonal periodicity (for details, see Supplementary Fig. 3 or reference 21). We observed that all the hexagonal autocorrelation peaks of individual maps exhibited a significance level of *P* < 0.01. In the average plot of autocorrelations, we confirmed that the peaks closely matched the expected hexagonal periodicity in three types of functional maps from species as different as cat, ferret, and galago (**Fig. 5e**).

## Discussion

How are various functional maps created? Is there a universal principle on the development of these functional architectures in the cortex? With these questions in mind, we introduced a novel model assuming that all the observed functional maps arise from projection of the structure of retinal mosaics (**Fig. 1**), and that this common origin induces systematic organization among the layouts of different maps, such as the orthogonal intersection of map gradients. In addition, we found hexagonal arrangement of iso-domains in each functional map by analysing the published animal data for various species, which has never been reported previously. These results support our idea of a single universal principle active in the developmental mechanism of diverse functional maps in the V1.

Several important issues on the development of functional maps were also addressed by our model. It has been reported that the ocular dominance map arises even before the inputs from both eyes are segregated in the cortex^23,40^, and this observation is understandable because the initial arrangement of ocular dominance domains does not require inputs from both eyes in our model. Considering the earlier development of the contralateral visual pathways^33^, our model hypothesis that spatial variation of inputs from the contralateral retinal mosaics is sufficient to induce the arrangement of the contralateral dominant domains (**Fig. 3d**) seems to be biologically plausible and is consistent with earlier observations of ocular dominance maps. The ipsilateral afferents are weak at this stage, so the cortical response is strongly modulated by the contralateral eye during early development^23^. Because the layout of an ocular dominance map can be determined by the input from one eye, the spatial relationship between ocular dominance and the orientation map is not ruined under monocular deprivation^41^.

The observed change in preferred frequency throughout the developmental stages is also explainable by the model. The V1 neurons show preference for relatively lower spatial frequency at the early stage of postnatal development, but preference tuning for higher spatial frequency arises at later stages^42^. Our model explains that this is because the selectivity for higher spatial frequency can develop when inputs from both eyes are provided to V1 neurons^9^. The anatomical connections of ipsilateral pathways arrive initially at V1 weeks before the observation of tuning for high frequency^40^. Effective connections are likely to take several weeks to be refined to match the orientation preferences between the two eyes for the complete development of the circuits^23^.

Compared to other maps, a relatively complicated architecture for spatial frequency maps may have caused difficulty in precise analysis of the location of orientation pinwheels on spatial frequency maps and resulted in contradictory results across some observations. Once, it was reported that pinwheels are preferentially located around the centre of ocular dominance domains^28^ where binocularity is relatively low. Because it was observed that the binocularity in the cortical neurons has positive correlation with the preferred spatial frequency across the cortical surface^9^, pinwheels were expected to be located in regions encoding low spatial frequency. However, in other, more precise optical imaging studies, the orientation pinwheels are reported to be more probably located at either low or high spatial frequency domains^7,25^ or in the positions between high- and low-frequency regions^43^. Considering that estimating the pinwheel location can be affected by the preprocessing condition found in the optical imaging methods (see Figure S2 in earlier work^44^), further studies imaging larger patches with higher resolution could provide a clue to resolve this contradiction.

On the other hand, one might question why neural tuning is arranged randomly in rodent V1, termed a salt-and-pepper map, and if our model can also address these issues in rodents. Recent tracing studies revealed that feature preference may not be randomly distributed in rodent V1, but is spatially clustered within a short distance that could not be observed with older techniques^45,46^. Apart from the distinct retinal organization found in higher mammals and rodents, another crucial factor in determining the scale of spatial clustering may be convergent projection in the feedforward visual pathway^39^. Compared to the localized convergence (almost one-to-one) of retinogeniculate connections in higher mammals^47^, the network of retinogeniculate connection is much more complex in rodents, so that a single LGN neuron receives input from a number of different types of RGCs^48^. Additional tracing experiments may reveal structural differences in the visual pathways of rodents and higher mammals and explain the origin of the diversity of functional maps across species.

The present scope of our model is limited to explaining how the retinal inputs could provide the “original” blueprint for each functional map. Obviously, the refinement of the circuits from cortical activity and visual experience is also a critical factor modifying the layout of functional maps^23^. To understand fully the development of the cortical circuits, it must be further investigated which component of the cortical circuit activity, such as cross-orientation suppression^49^, plays the most important role in each visual processing function.

Overall, our results suggest that the structure of retinal mosaics may provide an initial framework for multiple functional maps. This common origin may enable the development of various topographical correlations between different functional maps.

## Methods

The simulations were performed based on the statistical wiring model^18,19^. Here we briefly summarize the algorithm and the parameters used in the simulations.

### Structure of retinal ganglion cell mosaics

To simulate the development of binocular afferents to the V1, the contra- and ipsilateral RGC mosaics were individually generated by the superposition of ON and OFF RGC mosaics. The mosaic of each type of RGC was generated by adding random displacement to each vertex of a hexagonal lattice that represents the position of receptive fields. The centres of RGC receptive field position vectors are defined by

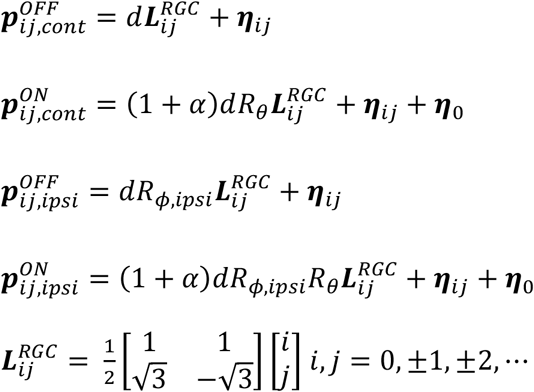

Here, 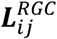 is the vertices of a unit hexagonal grid where *d* represents the grid spacing for the OFF mosaic, and (1 + *α*)*d* represents the grid spacing for the ON mosaic. The value of *d*, *α* was collected from published reconstruction data of RGC receptive fields of *M. fascicularis*^50^. The ***η**_ij_*, a random positional noise of RGC mosaics, is modelled as two-dimensional Gaussian noise to have a zero mean value with standard deviation *σ*_*n*_. To generate realistic RGC mosaics, *σ*_*n*_ was set to match the conditions of mean nearest-neighbour distance and noisiness observed in macaque monkey retinal mosaics^34,50^. A random relative spatial shift between ON and OFF mosaics ***η***_0_ was also added.

The matrix

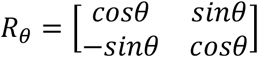

represents the relative rotation between the ON and OFF mosaics where *θ* is set to satisfy a scale factor proper for regenerating the actual period of orientation maps observed in macaque monkeys^51,52^. The matrix

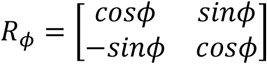

represents relative rotation between contralateral and ipsilateral RGC mosaics, where *ϕ* is a random value between 0 and *π*. All the parameter details are shown in Table 1.

**Table 1.**

| Parameters | Letter | Value |
| --- | --- | --- |
| Nearest OFF RGC distance | $d$ | $142\ \mu m^{50}$ |
| ON-OFF scaling factor | $\alpha$ | $0.105^{50}$ |
| Standard deviation of RGC positional noise | $\sigma_n$ | $17\ \mu m^{21,56}$ |
| Standard deviation of RGC to Cortical connection | $\sigma_{con}$ | $60\ \mu m$ |
| Standard deviation of retinotopic noise | $\sigma_{R\cdot}$ | $71\ \mu m$ |
| Nonlinearity of sigmoidal kernel | $\delta_{RGC}$ | 0.1 |
| Nonlinearity of sigmoidal kernel | $\delta_{V1}$ | 0.15 |
| Resource limit of single connection | $w_{lim}$ | 0.15 |
| Resource limit of total connection | $W_{lim}$ | 1 |
| Time constant of firing rate average | $\tau$ | 10 frame |
| Standard deviation of Gaussian kernel used in LHI calculation | $\sigma_{LHI}$ | $28\ \mu m$ |
| Learning rate in Hebbian learning | $\epsilon$ | $1 \times 10^{-5}$ |

### Initial conditions for connectivity parameters and receptive field models

At the initial stage of development, we assumed that the RGCs are statistically wired to cortical space with two-dimensional Gaussian function with a standard deviation of *σ*_*con*_.

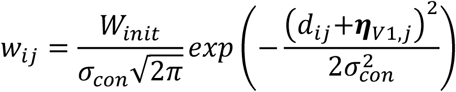

Here, *w*_*i*_ represents the synaptic weighting between i^th^ RGC and j^th^ cortical sites, where *W*_*init*_ is the initial connection weight. For realistic retinotopic projection, we added zero mean two-dimensional Gaussian noise, ***η***_*v1*_, with a standard deviation of *σ*_*R*_.

The receptive fields of RGCs and V1 neurons were defined as a centre-surround model of 2D Gaussian and their linear sum, respectively. The standard deviation of the surrounding region was set to three times that of the centre region^53^.

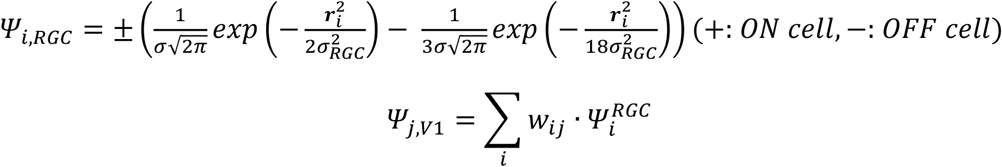

Here, *Ψ*_*i,RGC*_ is the receptive field of *i*^*th*^ RGC, and *Ψ*_*j,V1*_ is for the *j*^*th*^ cortical site where ***r***_*i*_ is a distance vector from the centre of *i*^*th*^ RGC to each position of the visual field. In this case, *σ*_*ON,RGC*_ and *σ*_*OFF,RGC*_ were set as (1 + *α*)*d*/2 and *d*/2 to satisfy the condition that the receptive fields of ON and OFF RGC mosaics cover all of the visual field^54^.

### Nonlinear response curve of a single neuron

The response curves of RGCs and V1 neurons were designed based on the linear-nonlinear model, which shows a normalized response to visual input ***S*** through a sigmoidal kernel

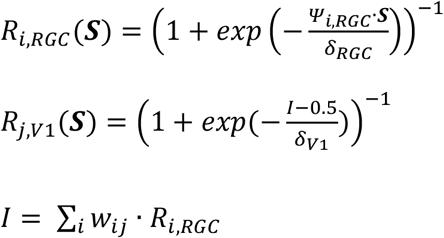

where *R*_*i,RGC*_(***S***) and *R*_*J,V1*_(***S***) are the response of i^th^ RGC and j^th^ cortical cell for visual stimuli ***S***. Here, *δ* stands for nonlinearity of the sigmoidal response function. All the parameter details are shown in Table 1.

### Binocular development and Hebbian plasticity

In our binocular development model, the development of connections between contralateral RGCs and V1 neurons were first simulated, and then the ipsilateral connections were allowed to develop. During development, drifting grating was given to binocular RGC mosaics where the drifting grating was designed as a sine function of twenty different spatial frequencies (0.5-8 cycles/degree) and twenty orientations (0-π). The synaptic weights were updated following a simple covariance rule^55^ as follows

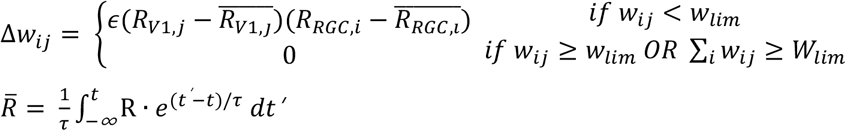

where the learning threshold *R̄* is defined as the average value of the current response of the cortical cell. The *τ* represents a time constant showing how rapidly the threshold changes. The *ϵ* represents the learning rate, how fast the synaptic weights are updated. Note that we assumed that there is a limitation of resource, so both *w*_*ij*_ and Σ_*i*_ *w*_*ij*_ have upper limit *w*_*lim*_ and *W*_*lim*_. The development of both contra- and ipsilateral pathways were simulated with drifting grating inputs for 10,000 frames.

### Measurement of cortical functional maps

At the initial stage of development, we assumed that the RGCs are statistically wired to cortical space in a two-dimensional Gaussian function with a standard deviation of ***σ***_***con***_.

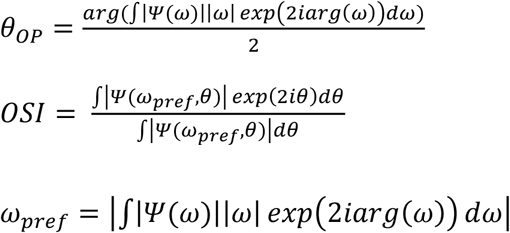

The preferred direction *θ*_*DP*_ of each cortical site was calculated from the angle between the centre positions of ON and OFF RGCs.

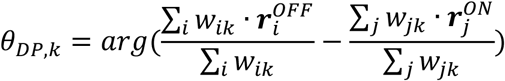

The ocular dominance was calculated as the relative strength of the mean cortical response 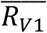, to visual stimulus *S*, given to contra- and ipsilateral RGCs.

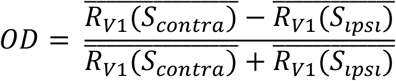

To measure the spatial frequency, we computed the response of cortical cells to drifting gratings of various orientation and spatial frequency. The preferred spatial frequency was defined as the value of spatial frequency that induces the maximum response. To implement a realistic map-topography, each map was simulated from the hexagonal RGC mosaics with a realistic level of spatial noise in the cell position^21^ and all the simulated maps in the study (except in Figure 2g–h) were averaged from 20 repeated trials of simulation.

### Analysis of cortical functional maps

To measure the intersection angle between map gradients, each functional map was first smoothed using a 2D Gaussian filter (*σ* = *d*/5). The gradients of OD and SF maps were calculated from the differences between the corresponding pixels of each map. The gradients of OP and DP maps were calculated in the complex domain: *C*(*OP*) = *exp*(2*i* ∗ *OP*) and *C*(*DP*) = *exp*(*i* ∗ *DP*). The intersection angle between maps was properly scaled into the [0, π] domain.

The local homogeneity index (LHI) measures how fast the orientation or direction changes at a given site, and was defined as the sum of complex vectors of OP or DP. In this calculation, a 2D Gaussian weighting of cortical distance was applied^49^.

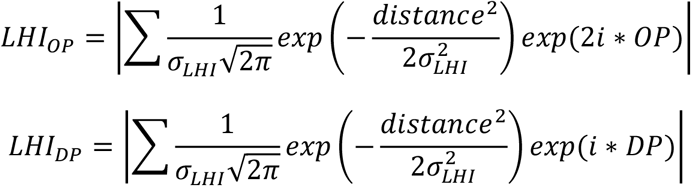

The overlap ratio between direction flip area and iso-orientation area was calculated as the ratio of overlapped regions to the entire area of iso-orientation domains. A direction flip was defined as the regions having the bottom 5% of LHI_DP_ and iso-orientation domain was defined as the regions having the top 5% of LHIop.

### Experimental data of functional maps and multi-electrode recordings

Multi-electrode data recorded from cats were provided by Jose-Manuel Alonso (ref. 15 and personal communication). The functional maps in the cat are from ref. 7, 10, 25, 27, 28 and 37 (orientation map), ref. 27 (direction map), ref. 7, 10, 25, 28 and 37 (ocular dominance map), and ref. 7, 10 and 25 (spatial frequency map). The functional maps in the monkey are from ref. 8, 9 and 38 (orientation map), ref. 9 and 38 (ocular dominance map) and ref. 8 and 9 (spatial frequency map). Orientation and ocular dominance maps in the galago are from ref. 26. Direction maps in ferret are from ref. 29 and 30.

## Acknowledgments

We are grateful to Jose-Manuel Alonso (State University of New York) for sharing receptive field data on the cat primary visual cortex. This research was supported by the Basic Science Research Program through the National Research Foundation of Korea (NRF), funded by the Ministry of Science and ICT (NRF-2016R1C1B2016039, NRF- 2016R1E1A2A01939949) (to S.P.).

## Author contributions

S.P. conceived the study. M.S., J.J. and S.P. designed the model. M.S. and J.J. performed the simulations. M.S., J.J., G.K. and S.P. analysed the data. M.S., J.J. and S.P. wrote the manuscript. All authors discussed and commented on the manuscript.

## Supplementary material

**Supplementary Figure 1.**
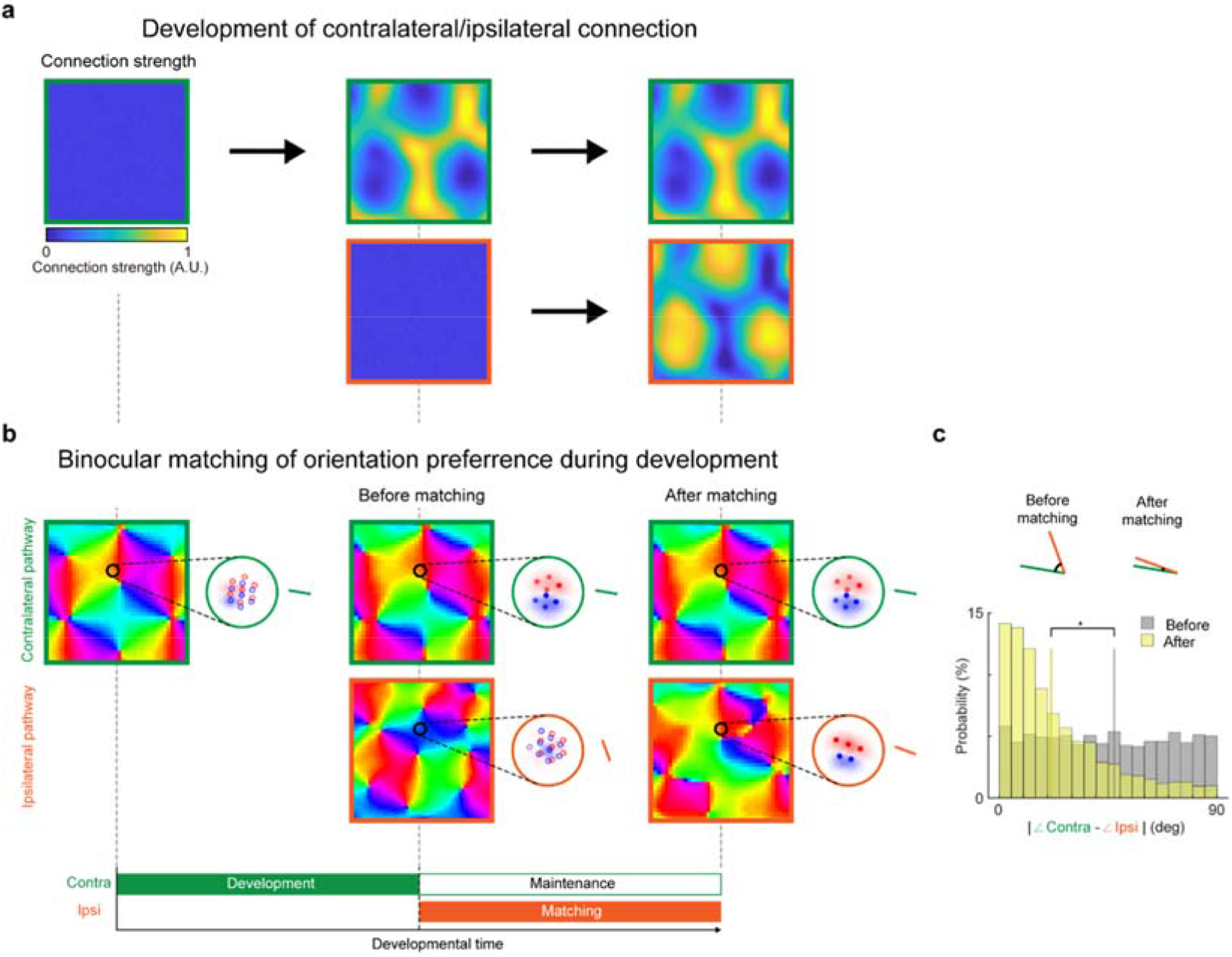
Orientation matching between contra- and ipsilateral pathways. **a**, Change in the strength of connections from contra- and ipsilateral pathways to V1 during development. (Left) Initially, only contralateral connections are weakly connected to the V1. (Middle) The contralateral connections are selectively strengthened by activity of postsynaptic V1 neurons. (Right) The ipsilateral connections are set up differentially, according to the connection strength from the contralateral pathway. **b**, Matching process of receptive fields and orientation tuning during the development of contra-and ipsilateral wirings. In the contralateral pathway, the RGC mosaic initially seeds the cortical map structure of orientation preference. In the ipsilateral pathway, the initial map is seeded by the ipsilateral RGC mosaics, independent of contralateral map structure. Next, during the matching process by activity-dependent plasticity, the connections from ipsilateral pathways are refined to match the orientation preference of the contralateral map^23^. **c**, Difference of preferred orientations between the two pathways before and after binocular matching.

**Supplementary Figure 2.**
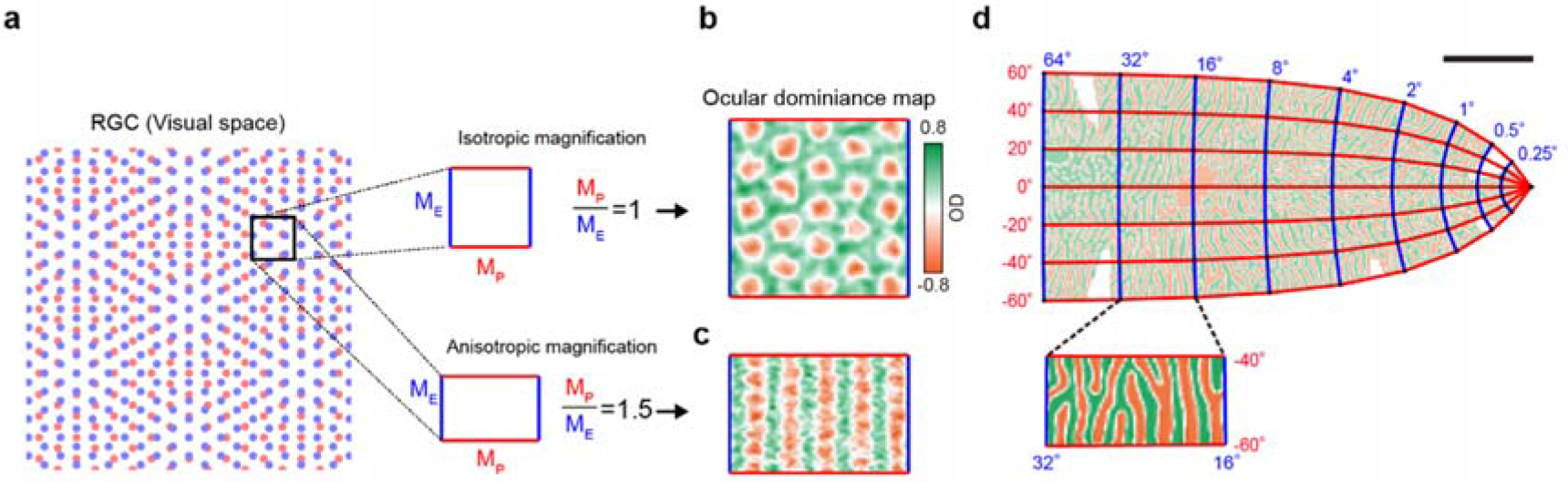
Stripe pattern of ocular dominance can be developed from moiré interference. **a**, Isotropic and anisotropic magnification factor observed in different species^34,35^. **b-c**, When anisotropy in retinotopy is significant (as in monkeys), the ocular dominance map is distorted so that the blobs of monocular regions are merged into stripe patterns of ocular dominance. **d**, As in our model simulation, in **c**, the stripe pattern of ocular dominance appeared to be parallel to the direction of shorter magnification, which is consistent with the direction of iso-eccentricity in monkeys^1^.

**Supplementary Figure 3.**
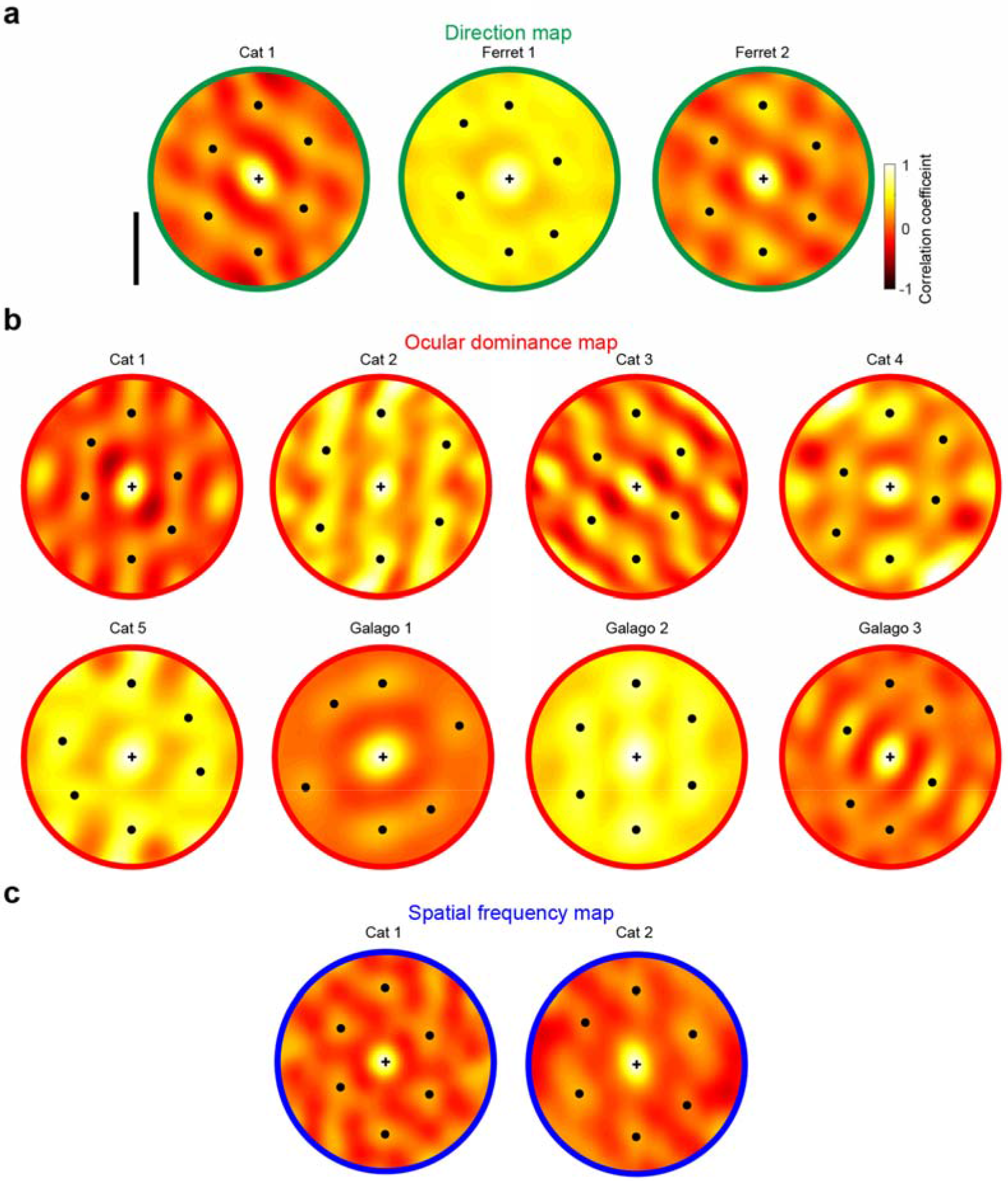
Hexagonal pattern of individual functional maps. Hexagonal pattern observed in the autocorrelation of individual functional maps. The magnitudes of local maxima (solid black dots) are statistically significant (bootstrap analysis, *P* < 0.01). **a**, Direction maps in cats^27^ and ferrets^29,30^. **b**, Ocular dominance maps in cats^7,10,25,28^ and galagos^26^, **c**, Spatial frequency maps in cats^7,25^.

